# MARTS-DB: A Database of Mechanisms And Reactions of Terpene Synthases

**DOI:** 10.1101/2025.05.11.653183

**Authors:** Martin Engst, Martin Brokeš, Tereza Čalounová, Raman Samusevich, Roman Bushuiev, Anton Bushuiev, Ratthachat Chatpatanasiri, Adéla Tajovská, Safa Mert Akmeşe, Milana Perković, Matouš Soldát, Josef Sivic, Tomáš Pluskal

## Abstract

**Background:** Terpene synthases (TPSs) are enzymes that catalyze some of the most complex reactions in nature – the cyclizations of terpenes, which form the carbon backbones to the largest group of natural products, the terpenoids. On average, more than half of the carbon atoms in a terpene scaffold undergo a change in connectivity or configuration during these enzymatic cascades. Understanding TPS reaction mechanisms remains challenging, often requiring intricate computational modeling and isotopic labelling studies. Moreover, the relationship between TPS sequence and catalytic function is difficult to decipher, while data-driven approaches remain limited due to a lack of comprehensive, high-quality data sources.

**Main:** We introduce the Mechanisms And Reactions of Terpene Synthases DataBase (MARTS-DB) – a manually curated, structured, and searchable database that integrates TPS enzymes, the terpenes they produce, and their detailed reaction mechanisms. MARTS-DB includes over 2,600 reactions catalyzed by 1,334 annotated enzymes from across all domains of life, with reaction mechanisms mapped as stepwise cascades for more than 400 terpenes. Accessible at https://www.marts-db.org, the database provides advanced search functionality and supports full dataset downloads in machine-readable formats. It also encourages community contributions to promote continuous growth.

**Conclusion:** User-friendly and comprehensive, MARTS-DB enables the systematic exploration of TPS catalysis, opening new avenues for computational analysis and machine learning, as recently demonstrated in the prediction of novel TPSs.

## Background

Terpene synthases (TPSs) are enzymes that catalyze some of the most complex reactions in nature – the cyclizations of terpenes. Compounds of diverse structure and function, terpenes are most notable for being the precursors to terpenoids, the largest class of natural products, featuring more than 100,000 known compounds [1, 2]. Their functions in nature are as varied as their structures, ranging from specialized plant metabolites to insect pheromones and steroids [3–6]. This remarkable chemical diversity originates from a relatively small pool of linear isoprenoid diphosphate precursors composed of connected five-carbon (C_5_) isoprene units and a diphosphate group [1, 7]. The biosynthesis of these precursors is performed by isoprenyl diphosphate synthases (IDSs), enzymes that are structurally similar to TPSs [1, 7, 8].

TPS reactions are initiated by the formation of a reactive carbocation through either diphosphate abstraction (class I TPSs) or alkene/epoxide protonation (class II TPSs). The active-site environment then guides the carbocation through a sequence of transformations, including cyclizations, hydride or methyl shifts, and proton transfers, concluding with deprotonation or hydroxylation [2, 9]. The precise sequences of steps in TPS reaction cascades have long been of interest to researchers. Isotope labelling experiments with precursor molecules have been routinely employed to trace atomic rearrangements within carbocations [10–14]. These insights are being further strengthened by quantum-chemical computations [15, 16]. In addition, computational structural studies and experiments with mutant TPSs all contribute to a deeper understanding of these processes [17, 18]. Although mechanisms for hundreds of terpene products have been described, no unified, publicly available resource systematically maps them.

The success of machine learning (ML) in biology, exemplified by protein language models [19, 20] and structure prediction tools like AlphaFold [21, 22], is fundamentally dependent on high-quality, well-annotated datasets. However, the power of ML for enzyme discovery and design remains largely untapped [23]. This is not due to a lack of powerful algorithms, but a scarcity of suitable data. Many state-of-the-art ML methods require a detailed mechanistic understanding of enzymatic reactions [19, 24, 25]. Consequently, machine learning cannot rely solely on large-scale databases such as UniProt or BRENDA, which do not primarily focus on reliably characterized enzymes [26, 27]. Specialized resources are therefore of great value to both data-driven studies and their respective research fields. For example, curated datasets exist for cytochromes P450 [28–30] or G protein-coupled receptors [31]. Terokit, a resource dedicated to terpenoid metabolism as a whole, has also been established [32]. However, a curated dataset of experimentally characterized TPSs is still lacking, and currently there is no resource that provides a unified, machine-readable map of TPS reaction mechanisms [28, 33].

To address this shortcoming, we present MARTS-DB (Mechanisms And Reactions of Terpene Synthases DataBase) – a manually curated, continuously updated database of characterized TPSs and their detailed reaction mechanisms. MARTS-DB is accessible through a user-friendly web interface that supports community data contributions. Though designed for a wide range of applications, MARTS-DB is particularly suitable for data-driven machine learning studies.

### Construction and content

At the time of MARTS-DB’s conception, the largest TPS dataset had been compiled by Durairaj et al., comprising 262 plant sesquiterpene synthases with major products [34, 35]. We expanded this dataset to encompass enzymes from all domains of life and different terpene types, while also extending the scope to incorporate minor products, cross-references, and detailed reaction mechanisms. We have mined TPS sequences from UniProtKB/Swiss-Prot and other available resources, based on their matches to TPS-specific Pfam domains (PF01397, PF03936, PF19086, PF13249, PF13243) [26, 36, 37]. The resulting dataset was further extended to include hundreds of manually curated enzymes from the published literature. Designated as MARTS-DB, the dataset currently includes 1,334 characterized TPSs and IDSs, which collectively catalyze the biosynthesis of more than 500 terpenes along with over 450 annotated reaction mechanisms.

Curation was performed by manually reviewing the primary literature for each recorded TPS. Only enzymes with experimentally proven activity and fully elucidated function were retained. The resulting dataset was automatically validated for errors related to the biochemical logic of terpene biosynthesis, such as substrate–product type consistency and reaction-type annotation. All manual annotations were systematically checked for consistency across the dataset. In addition, the dataset has been used and further validated by machine learning researchers. To keep pace with the continually evolving information on TPS reaction mechanisms, MARTS-DB records the version of each mechanism currently best supported by evidence.

Enzymes in MARTS-DB span all domains of life, including rare examples from viruses and archaea [38, 39]. Approximately two-thirds of the dataset are plant-derived TPSs, followed by fungal and bacterial sequences (**Fig. 1**) [40].

**Fig. 1.**
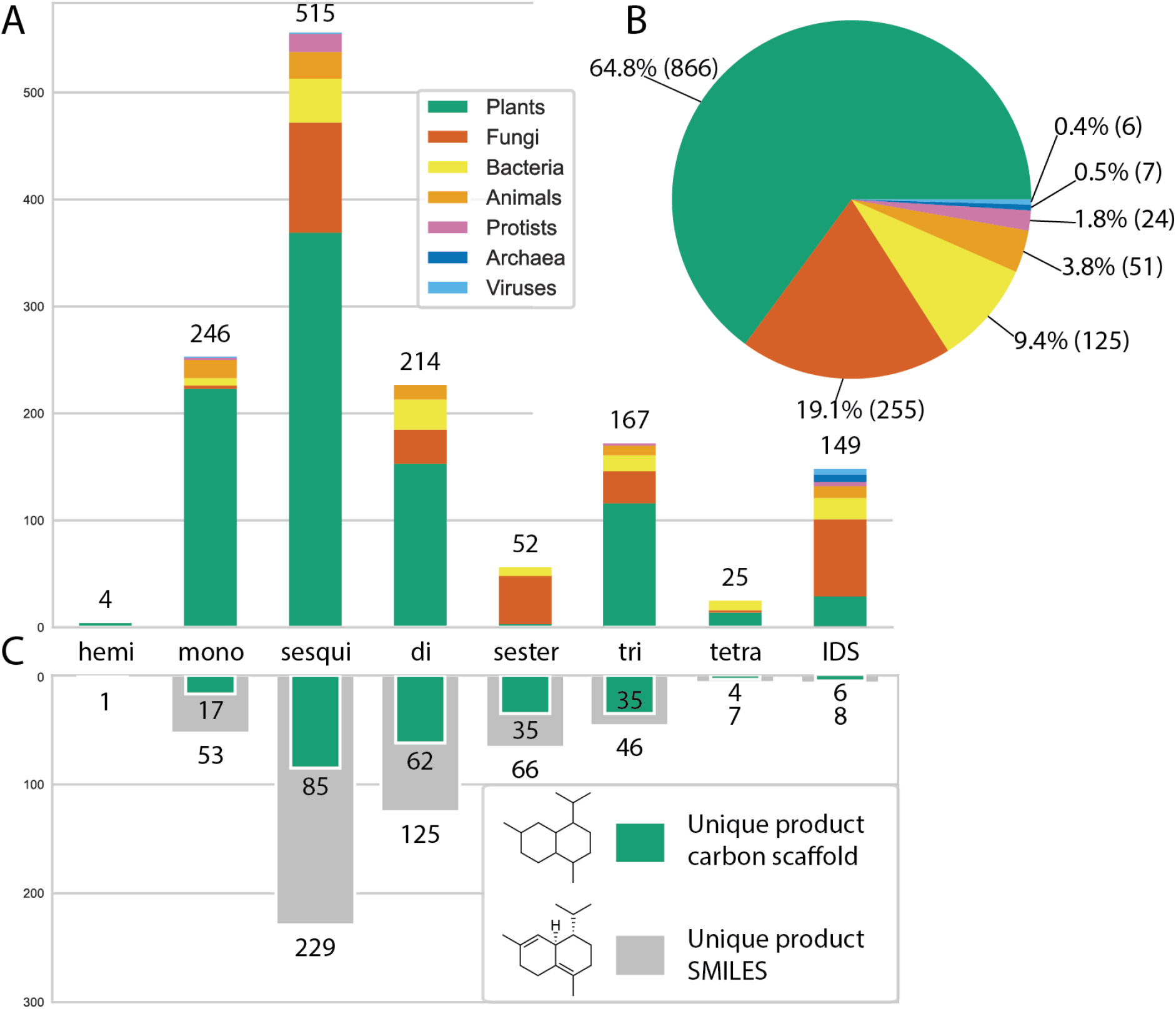
Number of terpenes and terpene synthases by type and taxonomic kingdom. (**A**) Number of TPSs catalyzing the production of each terpene type, coloured according to the taxon of their source species. TPSs catalyzing reactions of multiple terpene types are counted in each relevant category. (**B**) Distribution of taxonomic kingdoms for all 1,334 enzymes in MARTS-DB (data from August 2025). (**C**) Number of unique terpene products and unique carbon scaffolds of those products, grouped by terpene type.

Terpenes are generally categorized into types based on the size of their carbon backbone: hemiterpenes (C_5_), monoterpenes (C_10_), sesquiterpenes (C_15_), diterpenes (C_20_), sesterterpenes (C_25_), triterpenes (C_30_), and larger structures [1, 7]. Sesquiterpenes are the most represented group in MARTS-DB, followed by diterpenes, sesterterpenes, monoterpenes, and triterpenes. Even when considering only carbon scaffolds (ignoring stereochemistry and heteroatoms), sesquiterpenes remain the most abundant. Correspondingly, sesquiterpene synthases (sesquiTPSs) are the most common enzyme type in MARTS-DB, followed by monoterpene synthases (monoTPSs), and diterpene synthases (diTPSs). This distribution reflects the available literature and likely the apparent prevalence of sesquiterpenes in plants. However, as this distribution may also highlight a bias in the selection of the enzymes studied, users should be aware of this potential influence, particularly in machine learning applications [41, 42].

#### Database schema of MARTS-DB

MARTS-DB is a mySQL database structured around three core entities: Enzymes, Molecules, and Reaction Mechanisms. Each entity has a unique, persistent identifier (MARTS ID) in the format: marts_[EMR] followed by a five-digit number (E = Enzyme, M = Molecule, R = Reaction Mechanism). The five-digit number is assigned arbitrarily and does not encode any information about the entity. Even if a record is removed, its identifier can still be found and referenced (**Fig. 2**).

**Fig. 2:**
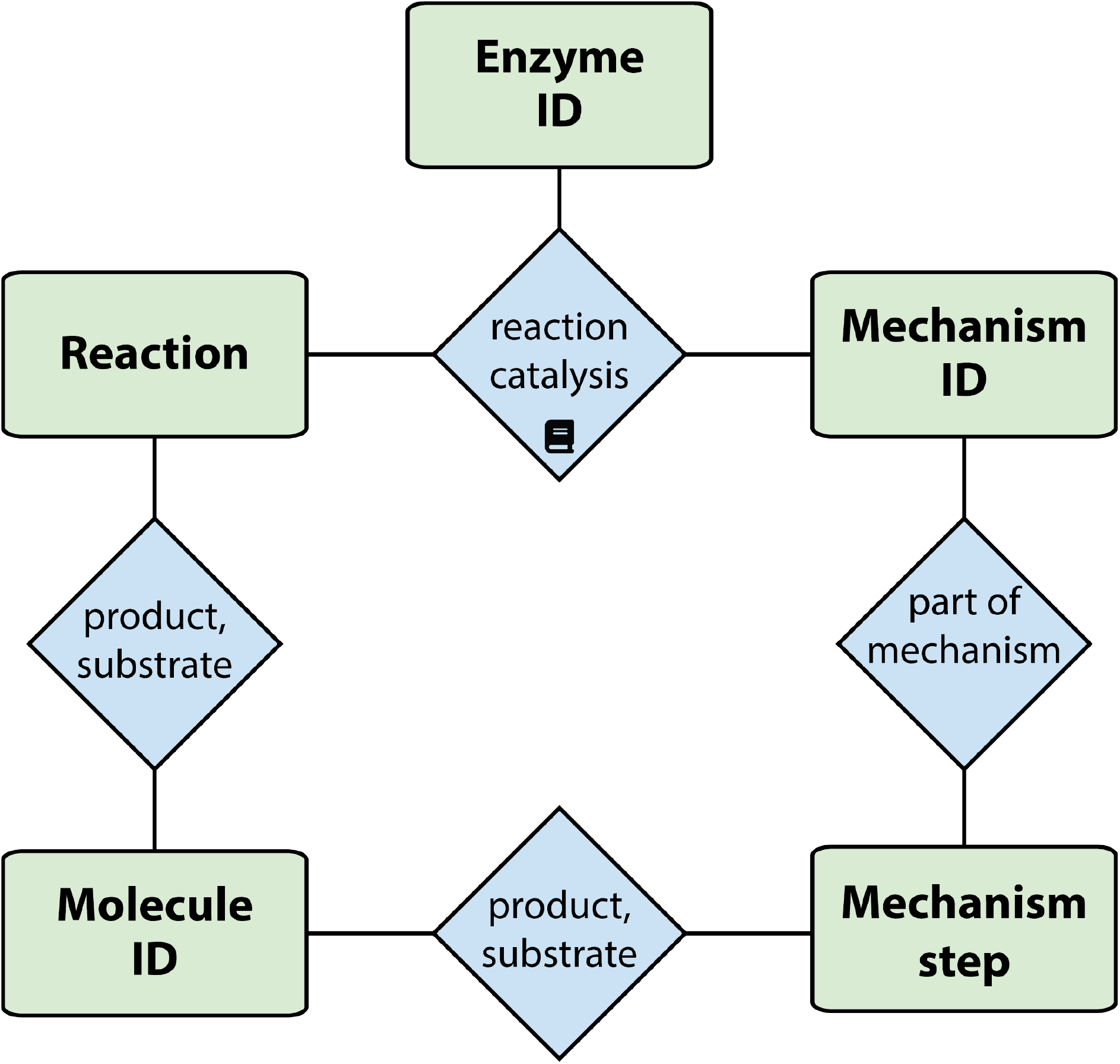
The schematic structure of MARTS-DB: The core of the database are Enzymes, Molecules and Reaction mechanisms, which can all be found by their persistent identifier called MARTS ID. Enzymes catalyze Reactions, where Molecules serve as either products or substrates. Mechanisms are composed of steps, whereas molecules again serve as either “products” or “substrates” of those steps. Each Enzyme-Reaction pair can be assigned a specific Mechanism. All Reaction–Enzyme and Reaction-Enzyme-Mechanism associations in the database are supported by one or more references to the primary literature, shown here by a book icon.

Molecules in MARTS-DB include both terpenes and reaction intermediates. Reactions are linked to the Enzymes that catalyze them, with each connection cross-referenced to the original literature source. This ensures accurate attribution in cases where a TPS has been investigated in multiple studies. Mechanisms are represented as sequences of atomic steps, each describing a single bond change, and are also linked to their literature sources. Every reaction step is assigned 1 of 16 step types reported in the literature (**Table S1**). Additionally, Mechanisms are assigned a specific category of evidence, whether experimental, computational, or purely theoretical. Each Enzyme–Reaction pair may have an associated Mechanism [43, 44].

Every entity with a MARTS ID is also linked to a history log, which tracks all changes to the entry, ensuring consistency of data over time.

### Utility and discussion

#### Web interface

MARTS-DB is freely accessible at https://www.marts-db.org/. The Home page provides basic statistics and a quick search function, which queries all fields in the database (**Fig. 3**). Search results are displayed as enzyme cards, including information on catalyzed reactions, with each Reaction and Mechanism linked to its source publication. Records can also be searched directly using the relevant MARTS ID.

**Fig. 3.**
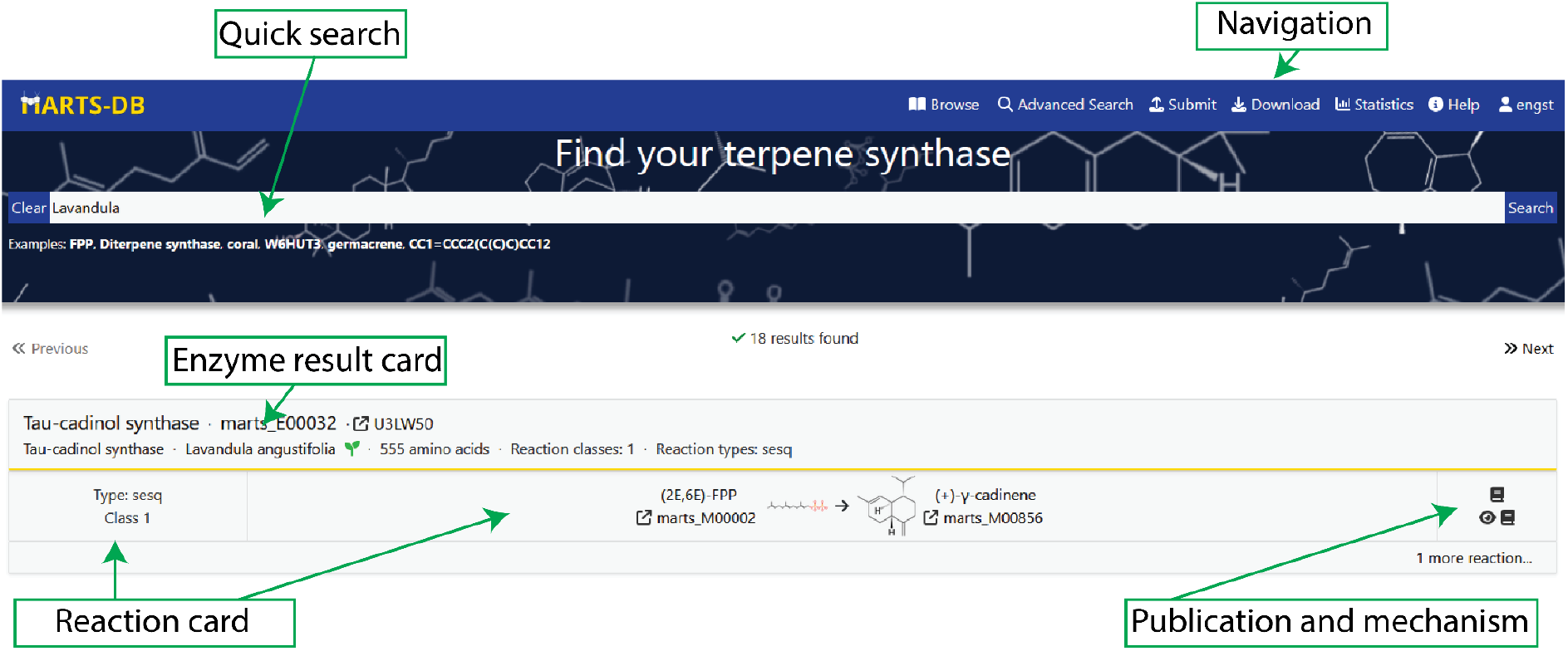
The MARTS-DB home page also serves as a quick search results page. Search results are displayed as cards for each enzyme, showing the reactions they catalyze.

The Browse pages enable users to explore the database by individual terpene products, Enzymes, or Reaction Mechanisms. These pages also function as dedicated search tools, providing extensive filtering options. Advanced search features support filtering by multiple parameters, including specific mechanism characteristics. A statistics page provides an overview of the database contents. Detailed documentation and usage instructions are available on the Help page.

Users can download the entire database through the dedicated Download page using the https protocol. This page provides not only the most recent version of MARTS-DB but also an archive of all previous versions. Archived versions are created approximately every two months, provided the data undergone substantial changes. This allows researchers to maintain backward compatibility in long-term computational studies, where data consistency across training and evaluation phases is essential. The database is distributed in CSV format, compatible with a wide range of analysis tools and programming environments. Details on the file format and its usage are described in the Help section. Additionally, users can download the entire network of reaction mechanism steps as a JSON file, which can be loaded in Cytoscape as well as all protein sequences in FASTA format [45].

Detailed Record pages are available for both Enzymes and Molecules. Enzyme records include taxonomic origin, reaction details (including associated mechanisms), and cross-references to UniProt and the National Center for Biotechnology Information (NCBI) [26, 46]. Conserved TPS motifs are highlighted in the sequence view. Molecule records include SMILES (including stereochemistry if assigned in the publication) and InChI identifiers, molecular properties, ChEBI cross-references (where available), and a list of associated Reactions and Enzymes, including detailed maps of Reaction mechanisms containing the molecule concerned. For both molecules and enzymes, the complete record history, linked to their MARTS ID, can also be accessed [26, 46, 47].

Mechanisms are presented as sequences of individual steps – viewable either in pop-up windows describing each step or as interactive diagrams within the Enzyme and Molecule Record pages as well as on a dedicated Mechanism browsing page. On the Browsing page, users can filter Mechanisms by terpene type, reaction class, the molecules involved, and the taxonomic origin of proteins capable of catalyzing the given Mechanism (**Fig. 4**).

**Fig. 4.**
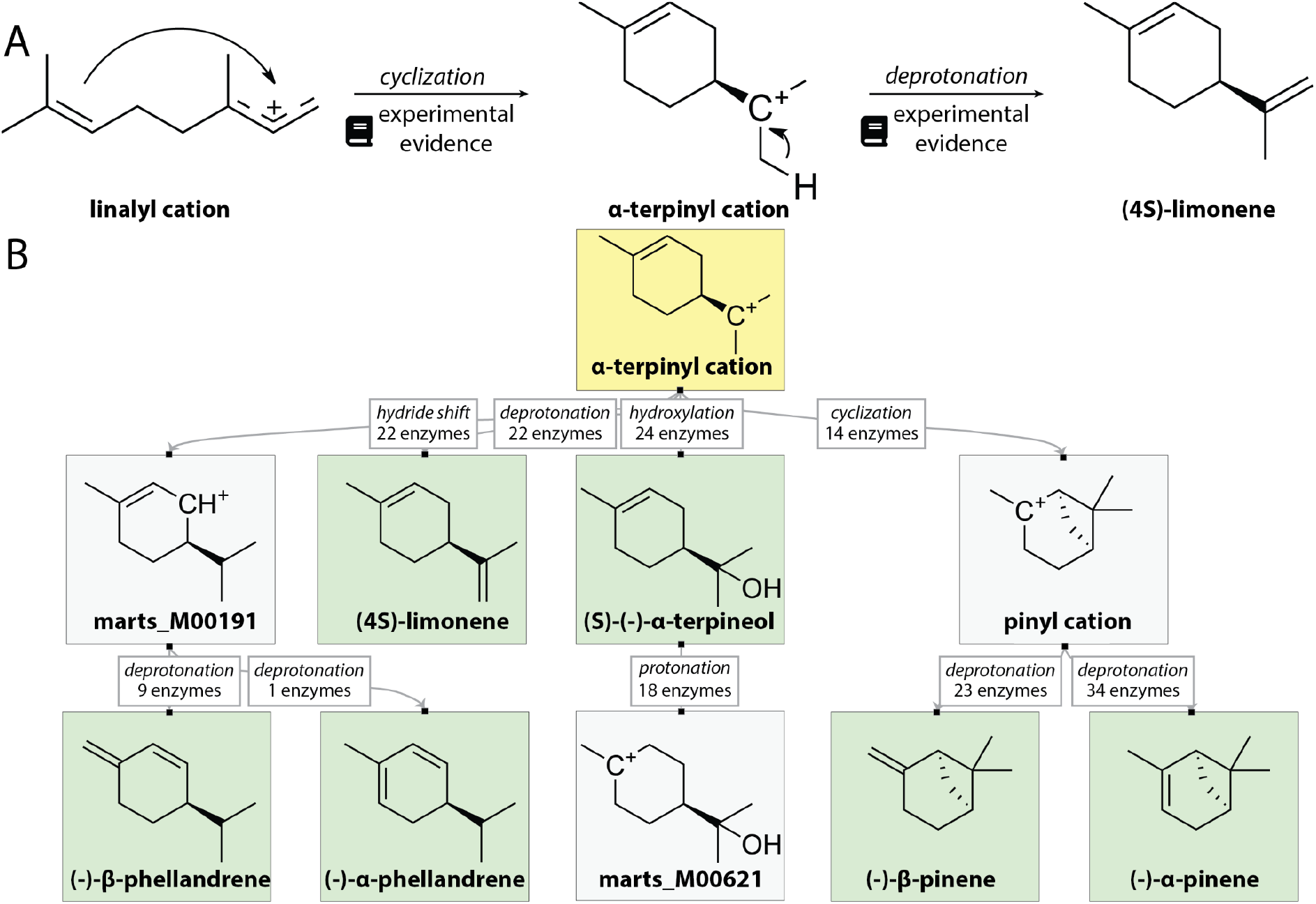
Mechanism representation in MARTS-DB. (**A**) Mechanisms are represented as individual steps of the reaction cascade. Each step approximates a change in a single bond. Each step is assigned a type and evidence sourced from the literature. (**B**) Example of the network representation of mechanisms within MARTS-DB web page. Final products are highlighted with green backgrounds, with yellow backgrounds indicating the molecular structure queried by the user.

Community contributions are enabled through a dedicated interface on the Submit page. Users can complete forms to add new enzymes and reactions or to assign additional reactions to existing enzymes. New mechanisms can also be defined for both newly added and existing reactions. To ensure consistency and appropriate attribution, the submit page is restricted to registered users; all submitted data undergo quality control before being incorporated into the database. This validation process follows the same principles applied to our data curation workflow, ensuring consistency with both the database structure and chemical logic

#### Future directions and utility

The TPS dataset contained in MARTS-DB has already been used as a training dataset for the EnzymeExplorer machine learning model designed to predict whether an unannotated enzyme is a TPS [48], as a hard test set for the enzyme screening tool CLIPZyme [49], and for evaluating the test-time model customization method ProteinTTT [50].

The dataset of TPS reaction mechanisms will also provide a valuable resource for future machine learning applications. Because terpene synthases exhibit a remarkable lack of sequence–function relationships, predicting the function of an unannotated enzyme remains a formidable challenge [51]. A promising approach may lie in the utilization of detailed mechanistic understanding, which would reframe the challenging biosynthetic product prediction task to a more tractable task of predicting a walk through the reaction mechanism tree.

Beyond serving as an annotated enzyme dataset for machine learning, MARTS-DB also provides a valuable resource for terpenoid researchers. For the first time, when the mechanism of a novel terpene synthase is elucidated, researchers can easily identify related mechanisms by querying the SMILES representations of the involved carbocations through the MARTS-DB Browse Mechanism interface. The dataset also opens new avenues of research into the evolutionary relationships of terpene synthases from a mechanistic perspective.

The database will continue to be curated and expanded by our team. Archived versions are expected to be added to the Download page approximately every two to four months, depending on the volume of new submissions. In addition to our curation, community contributions will be accepted. Long-term support for MARTS-DB is provided by the hosting institution (IOCB Prague) as well as current funding grants (see Funding section). Data integrity is maintained through regular backups to secure locations.

Additionally, we plan to expand MARTS-DB beyond its current focus on reaction mechanisms to include a comprehensive dataset of TPS structural data, a module devoted to mutational studies of TPSs, and a section dedicated to mass spectrometry-based detection of terpenes.

## Conclusion

MARTS-DB not only fills a gap in TPS data resources but also sets the stage for interdisciplinary research, offering a structured, searchable, and evolving platform that can be adapted to emerging analytical and computational approaches. Its meticulous curation and persistent identifier system ensure that researchers can trace every entry back to its source, maintaining the transparency and reproducibility required for robust scientific investigation.

As MARTS-DB continues to expand, it is expected to become an indispensable tool for the study of terpene biosynthesis, supporting the annotation of novel enzymes, the discovery of new natural products, and the development of predictive models for enzyme function. Finally, the community-driven aspect of the platform will ensure it remains up to date and relevant, reflecting the collective knowledge and ongoing contributions of researchers in the field.

## List of abbreviations

IDS: isoprenyl diphosphate synthase
MARTS-DB: Mechanisms and Reactions of Terpene Synthases Database
NCBI: National Center for Biotechnology Information
TPS: terpene synthase

## Declarations

### Ethics approval and consent to participate

Not applicable.

### Consent for publication

Not applicable.

### Availability of data and material

The MARTS-DB dataset is provided under the CC BY 4.0 license (https://creativecommons.org/licenses/by/4.0/) and can be freely downloaded from the database web page (https://www.marts-db.org). The current version of the dataset (as of October 2025) is published on Zenodo under the DOI https://doi.org/10.5281/zenodo.17313803.

### Authors’ contributions

T.P. conceptualized the project. M.E, A.T., T.Č., R.S., R.Ch., R.B, A.B, and M.P. collected and curated the dataset. S.M.A, R.B, A.B and M.S. validated the data. M.E. and M.B. created the web interface. M.E. wrote the manuscript. R.B., A.B., S.M.A and M.S. significantly contributed to the revision. T.P. and J.S. supervised the project. All authors approved the final manuscript.

## Acknowledgements

Not applicable

