## Supplementary material for "MARTS-DB: A Database of Mechanisms And Reactions of Terpene Synthases": Table S1

### Supplementary information

| Step type | Description |
| --- | --- |
| Dephosphorylation | Abstraction of the diphosphate group, the initiation step for class I TPS reactions [1, 2]. |
| Protonation-class II | Protonation of an alkene or epoxide bond, initiating a class II TPS reaction [9]. |
| Cyclization | Nucleophilic attack of a double bond on the positive charge of a carbocation, forming a new cycle within the molecule [2]. |
| Hydride shift | Transfer of a single hydride ion between a carbon atom and a carbocation [51]. |
| Methyl shift | Transfer of a methyl group between a carbon atom and a carbocation [1, 52, 53]. |
| WM rearrangement | Wagner–Meerwein rearrangement, typically a 1,2 migration of a ring carbon atom in a polycyclic system. In TPS literature and MARTS-DB, the term applies to any shift in the ring skeleton [54, 55]. |
| Proton transfer | Transfer of a proton between a hydrogen atom and a double bond, resulting in a repositioning of a double bond on the positive charge [56, 57]. |
| Bond cleavage | Cleavage of a C–C bond, resulting in the formation of a new double bond [58–60]. |
| Oxygen cyclization | Nucleophilic attack reaction of an OH group on a carbocation, abstracting the alcohol hydrogen and forming a heterocycle [61, 62]. |
| Phosphorylation | Re-addition of the abstracted pyrophosphate to the carbocation. Via phosphorylation, TPSs achieve isomerization of a *trans*-allylic cation to a *cis*-allylic cation, the first step in monoterpene and some sesquiterpene and diterpene biosynthesis [1, 63, 64]. |
| Protonation | Protonation of a stable intermediate formed during the reaction cascade [65–67]. |
| Deprotonation | Abstraction of a proton from the carbocation, forming a double bond [65, 68]. |
| Hydroxylation | Nucleophilic attack of water on a carbocation, forming an alcohol and ending the reaction cascade [18, 69]. |
| Cyclopropanation | Rare termination of the reaction cascade where cyclization |

| | |
|---|---|
| deprotonation | coupled with deprotonation forms a cyclopropyl ring [66, 70]. |
| Fragmentation | Observed in terpenes created from unusual precursors with non-canonical carbon numbers (for example, $C_{16}$), where the carbocation fragments into two parts [52, 71]. |
| Cycloaddition | Mending of a fragmented carbocation by cycloaddition [52]. |

**Table S1**. The 16 mechanism steps used in MARTS-DB to describe the reaction mechanisms of terpene synthases.
